# Partial Correlation Analysis Reveals Abnormal Retinotopically Organized Functional Connectivity of Visual Areas in Amblyopia

**DOI:** 10.1101/237362

**Authors:** J.D. Mendola, J. Lam, M. Rosenstein, L.B. Lewis, A. Shmuel

**Affiliations:** Departments of Ophthalmology, McGill University, Montreal, QC H9G 1A4; Neurology and Neurosurgery, McGill University, Montreal, QC H9G 1A4; Montreal Neurological Institute, McGill University, Montreal, QC H9G 1A4

**Keywords:** Functional MRI, spontaneous activity, intrinsic, resting state functional magnetic resonance imaging, resting state functional connectivity, partial correlation, regional, strabismus, binocular vision, suppression

## Abstract

Amblyopia is a prevalent developmental visual disorder of childhood that typically persists in adults. Due to altered visual experience during critical periods of youth, the structure and function of adult visual cortex is abnormal. In addition to substantial deficits shown with task-based fMRI, previous studies have used resting state measures to demonstrate altered long-range connectivity in amblyopia. This is the first study in amblyopia to analyze connectivity between regions of interest that are smaller than a single cortical area and to apply partial correlation analysis to reduce network effects. We specifically assess short-range connectivity between retinotopically defined regions of interest within the occipital lobe of 8 subjects with amblyopia and 7 subjects with normal vision (aged 19-45). The representations of visual areas V1, V2, and V3 within each of the four quadrants of visual space were further subdivided into three regions based on maps of visual field eccentricity. Connectivity between pairs of all nine regions of interest in each quadrant was tested via correlation and partial correlation for both groups. Only the tests of *partial* correlation, i.e., correlation between time courses of two regions following the regression of time courses from all other regions, yielded significant differences between resting state functional connectivity in amblyopic and normal subjects. Subjects with amblyopia showed significantly higher partial correlation between para-foveal and more eccentric representations within V1, and this effect associated with poor acuity of the worse eye. In addition, we observed reduced correlation in amblyopic subjects between isoeccentricity regions in V1 and V2, and separately, between such regions in V2 and V3. We conclude that partial correlation-based connectivity is altered in an eccentricity-dependent pattern in visual field maps of amblyopic patients. Moreover, results are consistent with known clinical and psychophysical vision loss. More broadly, this provides evidence that abnormal cortical adaptations to disease may be better isolated with tests of partial correlation connectivity than with the regular correlation techniques that are currently widely used.

## 1 Introduction

Amblyopia is the most common visual disorder in children, with a prevalence of 2-4% in the general population. Amblyopia is clinically defined as decreased acuity (typically 20/30-20/80) in an otherwise healthy eye. Impairments result not from ocular lesion but from abnormal neural development associated with atypical early visual experience. The most common etiological types of monocular amblyopia are those associated with strabismus, anisometropia, or both (von Noorden & Campos, 2001, McKee *et al.*, 2003; Joly & Frankò, 2014). Strabismus is the uncorrected misalignment of the visual axis, while anisometropia is a large refractive imbalance of the two eyes. Amblyopia is a syndrome which is often described by deficits in acuity, contrast sensitivity, and stereoscopic depth perception, but also represents impairments in form perception, spatial localization, fixation, accommodation, crowding, attention, motion perception, ocular motility, and temporal processing (e.g., Asper *et al.*, 2000; McKee *et al.*, 2003). Visual deficits in amblyopia are usually, but not always monocular (e.g., Hamm *et al.*, 2014; Hou *et al.*, 2016), and generally impact the high acuity of central vision to a greater extent than peripheral vision (Levi *et al.*, 1984; Kiorpes *et al.*, 1998; Hess & Pointer 1985; Sireteanu & Fronius, 1990; Levi & Walters, 1997; Conner *et al.*, 2007a; Babu *et al.*, 2013; Li *et al.*, 2017).

Based on numerous psychophysical, physiological (e.g., Levi *et al.*, 1984; Movshon *et al.*, 1987; Sengpiel & Blakemore, 1996; Kiorpes *et al.*, 1998) as well as neuroimaging (Mendola *et al.*, 2005; Conner *et al.*, 2007b; Ding *et al.*, 2013; Lin *et al.*, 2012; Joly & Frankò, 2014; Brown *et al.*, 2016) studies, the physiological basis for amblyopia is believed to begin at the level of primary visual cortex where binocular processing commences. This premise is consistent with widespread observations that abnormal suppressive interocular interactions characterize amblyopia (e.g., Sireteanu, 1982; Sengpiel & Blakemore, 1996; Agrawal *et al.*, 2006; Conner *et al.*, 2007b; Farivar *et al.*, 2011; Adams *et al.*, 2013; Hess *et al.*, 2014; Birch *et al.*, 2016), and the traditional clinical premise that pre-cortical sites are essentially normal. Nevertheless, some studies have reported altered morphology and function in the lateral geniculate nucleus (LGN) of amblyopic animal models and of amblyopic human subjects (e.g., Barnes *et al.*, 2010; Ding *et al.*, 2013; Joly & Frankò, 2014; Crewther & Crewther, 2015). In addition, fMRI studies have reported reduced LGN activity (Hess *et al.*, 2009b), and reduced Granger effective connectivity from LGN to V1 for amblyopic eye viewing conditions in six subjects of mixed etiology (Li *et al.*, 2011). Although LGN-V1 feedforward and feedback connectivities were similarly affected, only the forward effect showed an ipsilateral specificity. Therefore, a modest feedforward deficit is possible (e.g., Brown *et al.*, 2013), although these LGN losses are likely to also be influenced by altered modulatory feedback connections from V1. Overall, evidence suggests that deafferentation of cortical binocular cells in V1 is a primary site of abnormality that is possibly reinforced at the level of the LGN (Joly & Frankò, 2014).

Both strabismic and anisometropic amblyopia tend to show a selective perceptual loss of the central visual field that relies on the fovea (Levi *et al.*, 1984; Sireteanu & Fronius, 1990; Levi *et al.*, 2002; Hess & Pointer 1985; Movshon *et al.*, 1987; Kiorpes *et al.*, 1998; Asper *et al.*, 2000; Babu *et al.*, 2017). In addition, more recent neuroimaging studies have generally come to consistent conclusions (Muckli *et al.*, 2006; Conner *et al.*, 2007b; Clavagnier *et al.*, 2015; Huang *et al.*, 2017). Conner and colleagues (2007a) noted the recruitment of parafoveal representations in cortical positions in the occipital pole that represented foveal mapping in control eyes. They suggested that the susceptibility of foveal representations to interocular mismatch, and to compensatory interocular inhibition, is higher compared to other localized representations, due to the small receptive field sizes of central vision (e.g., Sireteanu, 1982). This concept is supported by past psychophysical studies that noted qualitative similarities between the amblyopia fovea and the normal periphery (Levi *et al.*, 1984).

In recent years, the value of resting state functional magnetic resonance imaging (rs-fMRI) has been demonstrated in the study of amblyopia (Lin *et al.*, 2012; Ding *et al.*, 2013; Wang *et al.*, 2014; Liang *et al.*, 2016). Resting state, unlike task-based fMRI, requires neither stimulation nor response and may serve as an indicator of intrinsic brain function. These signals represent the spontaneous neuronal activity and connectivity of the human brain (Fox & Raichle, 2007; Shmuel & Leopold, 2008; Smith *et al.*, 2009; Wang *et al.*, 2013). At rest, distant cortical areas that are functionally related often show coherent spontaneous, slow (<0.1 Hz) fluctuations in fMRI signal (Biswal *et al.*, 1995; Nir *et al.*, 2006). It has been suggested that the underlying anatomical connections shape the spatial pattern of resting-state functional connectivity (De Luca et al., 2006; Fox and Raichle, 2007; van den Heuvel et al., 2009; Wang et al., 2013), although functional connectivity is not identical to anatomical connectivity (Honey et al., 2010). The differences between structure and function can be attributed in part to the contribution of polysynaptic interactions to resting-state functional connectivity (Vincent et al., 2007; Shmuel and Leopold, 2008; Honey et al., 2009; Adachi et al., 2012).

A few studies have used rs-fMRI to evaluate whole-brain long- and shorter-range functional connections in children or adults with anisometropic amblyopia (Lin *et al*., 2012; Ding *et al*., 2013; Wang *et al*., 2014; Liang *et al*., 2016). Overall, abnormal patterns were reported for many brain regions including calcarine, middle occipital, precuneous, posterior parietal cortex, posterior frontal cortex, and cerebellum in amblyopic subjects. Presumably, these latter regions are involved in visuomotor action and visuospatial attention, among other functions, which appear to be impaired in amblyopes (Niechwiej-Szwedo *et al.*, 2012, 2014). Two similar studies have now examined subjects with strabismic amblyopia and also find multiple sites of abnormality including lingual and inferior temporal cortex, as well as cerebellum, angular and cingulate and medial frontal cortex (Huang *et al.*, 2016; Tan *et al.*, 2016). While these wide-ranging cortical effects are likely to result from the chronically impaired visual input experienced in visual cortex, none of these studies have actually defined the functional boundaries of individual visual areas or evaluated the connectivity between them.

Recent evidence suggests that correlations in rs-fMRI do in fact reflect functional organization at a scale finer than that of individual cortical areas (Dawson *et al*., 2016). For example, the synchronous fluctuations between topographically organized visual areas might be related to their shared intrinsic retinotopy, similar to the retinotopically-targeted anatomical connections known from monkey visual cortex (e.g., Amir *et al*., 1993; Angelucci *et al*., 2002; Shmuel *et al.*, 2005). Indeed, such eccentricity-specific correlations have been demonstrated between visual areas in normal macaques and humans (Vincent *et al.*, 2007; Heinzle *et al.*, 2011; Arcaro *et al*., 2015; Dawson *et al*., 2016).

In the current study, we assess how resting state functional connectivity in visual areas depends on the level of visual hierarchy and eccentricity in amblyopic as well as visually normal control subjects. We use methods that have been previously validated and applied to normal visual cortex (Dawson *et al.*, 2013; Dawson *et al.*, 2016). By specifically studying rs-fMRI signals in the first three visual areas, V1, V2, V3, our goal is to determine how hierarchical connectivity *between* retinotopic visual areas differs in amblyopia. Furthermore, we ask how the connectivity between sub-regions *within* V1-V3, representing different eccentricities of the visual field, might differ in amblyopia. Based on the evidence reviewed above, we hypothesize that cortical regions representing the central visual field have a greater degree of abnormality in organization compared to regions representing the peripheral visual field in amblyopic patients. Our aim is to specifically test the representations within the occipital lobe with the most relevance to visual acuity loss in central vision.

## 2 Materials and Methods

### 2.1 Subjects

We studied 15 adult volunteers aged 19 - 45 years (13 female, 2 male). Seven were control subjects with normal or corrected-to-normal vision, six had previously been diagnosed with strabismic amblyopia, and two had previously been diagnosed with anisometropic amblyopia. Any subject with other known or suspected neurological or psychiatric conditions was excluded. These subjects were recruited through public advertisement in Montreal. Informed consent was obtained from all subjects, in accordance with the Code of Ethics of the World Medical Association (Declaration of Helsinki).

Our subject groups were matched for mean age (control, 28.6 years, SD = 7.6; amblyopic, 27.9 years, SD = 8.9).9. All amblyopic subjects had a history of patch occlusion treatment during childhood, but the presence of visual impairment at the time of testing demonstrates that the deficit was never completely reversed. One of the six strabismic subjects also reported surgical correction of their deviation in childhood.

All subjects completed a screening to confirm their diagnosis. Diagnosis of anisometropic amblyopia was assigned on the basis of: (1) interocular refractive difference of hyperopia ≥ +1.0 diopter, astigmatism ≥ +1.0 diopter, or myopia ≥ −2.5 diopters; or (2) history of anisometropia but no history of strabismus or strabismus surgery. Diagnosis of strabismus was made on the basis of a history of strabismus or strabismus surgery, but no anisometropia (as defined above). In clinical practice it is common to find that some subjects with amblyopia present with a mixed anisometropic–strabismic diagnosis, although little consensus exists regarding additional subtypes. The direction and magnitude of strabismic deviation in our subjects was determined with cover–uncover, alternate cover and prism testing. Three of our strabismic subjects showed inward eye deviations (esotropia) and three of our strabismic subjects showed outward deviations (exotropia). Manifest deviations for our subjects ranged from 0 to 14 prism diopters. The ophthalmological tests also included examination of the fundus with dilation, documentation of any ductions and versions, autorefraction, and a sensory exam including Snellen visual acuity (Lombart Instrument, Norfolk, VA, USA), and stereoacuity (Titmus Optical, Inc., Petersburg, VA, USA). The degree of stereoacuity was measured with the Titmus stereoacuity test, scored according to highest level of detectable horizontal disparity for the Wirt rings or for the Titmus fly. The crudest stereoacuity measurable with this test is 3000 arcsec, assigned for patients able to perceive disparity only in the Titmus fly illustration. In addition to diagnosis validation, these results provide a general indication of the residual amount of binocular integration in each subject. See Table 1 for more information on each subject. All subjects completed 2 scan sessions: a scan session to obtain high-resolution anatomical images and resting state functional data, and a scan session for functional retinotopy data (2^nd^ scan session).

**Table 1.** Amblyopic Subjects*

|  |  |  |  |  |  |  | Eye Acuity (Snellen/logMAR) |  | Stereoaucuity |
| --- | --- | --- | --- | --- | --- | --- | --- | --- | --- |
| Subject | Gender | Age | Aniso-metropia | Esotropia | Exotropia | Deviation (degrees) | Fellow Eye | Amblyopic Eye |  |
| S1 | Female | 26 | No | No | **Yes** | 1 | 20/30/0.2 | 20/100/0.7 | 800" |
| S2 | Female | 45 | No | No | **Yes** | 8+5 HT | 20/20/0 | 20/160/0.9 | 3000" |
| S3 | Female | 25 | **Yes** | No | No | N/A | 20/16/-0.1 | 20/40/0.3 | 70" |
| S4 | Female | 22 | No | No | **Yes** | 10 | 20/25/0.1 | 20/40/0.3 | 3000" |
| S5 | Female | 23 | No | **Yes** | No | 14 | 20/20/0 | 20/32/0.22 | None |
| S6 | Female | 19 | No | **Yes** | No | 1 | 20/20/0 | 20/360/>1.0 | None |
| S7 | Female | 25 | No | **Yes** | No | 4 | 20/20/0 | 20/400/>1.0 | None |
| S8 | Female | 38 | **Yes** | No | No | N/A | 20/20/0 | 20/32/0.22 | 400" |

\*Shaded rows indicate the subjects with worse acuity scores (i.e., higher logMAR values), plotted in Fig. A with circles.

### 2.2 Resting state fMRI

Resting state data was collected during the 1^st^ scan session. During the resting state functional scans, the subjects kept their eyes closed. Subjects were scanned in a Siemens 3T Magnetom TrioTim scanner, using a 32-channel phased array head coil. Echo-planar imaging (EPI) was used to measure blood oxygenation level-dependent (BOLD) changes in image intensity. Each run consisted of 270 contiguous EPI whole-brain functional volumes [repetition time (TR) = 2000 ms; echo time (TE) = 30 ms; flip angle = 90°; 38 slices; matrix = 64 × 64; field of view (FOV) = 230 mm; acquisition voxel size = 3.6 × 3.6 × 3.6 mm].

In the same session as the resting state data acquisition, 3 high-resolution anatomical volumes were acquired and averaged together to improve the signal-to-noise ratio. For each of the 3 volumes, three-dimensional (3D) anatomical images were acquired via averaging together 3 consecutive T1-weighted MPRAGE (magnetization-prepared rapid gradient echo) sequences (312 s each), containing 176 interleaved 1.0 mm slices with 1.0 mm × 1.0 mm in-plane resolution, oriented along true AC-PC (TR = 2300 ms, TE = 3.4 ms, flip angle = 90°, FOV = 256 mm, voxel resolution = 1.0 mm × 1.0 mm × 1.0 mm).

Resting state runs were preprocessed using the FMRIB Software Library's FEAT software package (Jenkinson *et al.*, 2012). Using the pre-stats option, we discarded the first three volumes to avoid non steady-state effects, performed motion correction, high-pass filtering (0.01 Hz), slice timing correction, and registration to the T1-weighted (MPRAGE) anatomical volume from the retinotopy session. Spatial smoothing and co-registration between subjects were not applied to avoid unnecessary signal spread over space.

Following the FSL FEAT pre-stats step, the resting state functional runs underwent denoising. A cerebrospinal fluid (CSF) and white matter (WM) masks were created by segmenting the resting state anatomical volume using FSL's FAST software package (Zhang *et al.*, 2001). The masks were then eroded by one voxel, thresholded (80% tissue type probability), and registered to each preprocessed functional volume. The average time courses from the CSF and WM masks were obtained and the two time courses were regressed out of the functional volume. Since we analyzed resting state fMRI data from a relatively small part of the brain, neither global signal regression (Carbonell *et al.*, 2011) nor correction for the impact of the global signal on functional connectivity (Carbonell *et al.*, 2014) were performed.

### 2.3 Retinotopy

Retinotopy data was collected during the 2^nd^ scan session. Subjects were scanned in a Siemens 3T Magnetom TrioTim scanner, using a 20-channel phased array head coil. After a localizing scan was obtained, a high-resolution anatomical volume was pursued. For this volume, three-dimensional (3D) anatomical images were acquired using a T1-weighted MPRAGE (magnetization-prepared rapid gradient echo) sequence, containing 176 1.0 mm thick slices with 1.0 mm × 1.0 mm in-plane resolution (TR = 2300 ms, TE = 3.0 ms, flip angle = 9 deg, FOV = 256 mm, voxel resolution = 1.0 mm × 1.0 mm × 1.0 mm). This anatomical volume was later utilized to register functional retinotopy data from the 2^nd^ session to the cortical surface model obtained from the high-resolution anatomical data in the 1^st^ session. Retinotopy data was acquired with blood oxygenation level-dependent (BOLD) functional imaging, using 28 interleaved 3.0 mm thick slices with 2.1 mm × 2.1 mm in-plane resolution, oriented obliquely parallel to the calcarine sulcus (TR = 2000 ms, TE = 30 ms, flip angle = 90 deg, FOV = 263 mm, voxel resolution = 2.1 mm × 2.1 mm × 3.0 mm).

Retinotopy stimuli were projected onto a rear-projection screen visible from within the MRI scanner via an angled mirror. All subjects viewed the stimulus with both eyes open under natural conditions. The visual stimuli were projected from a liquid crystal display (LCD) projector at 1024 × 768 resolution and 60 Hz refresh rate onto a translucent screen at the end of the scanner bore. The subjects viewed the screen at the total viewing distance of 138 cm through a mirror mounted to the coil, which yielded 32° × 24° (40° in diagonal) of viewing angle. A central fixation target was presented at all times during retinotopy functional scans. Subjects were instructed to maintain fixation on this target throughout the scan. The target was a small red arrowhead (0.5°) pointing in one of four directions (i.e., up, down, left or right), which randomly changed direction every 4 s. In order to aid fixation stability and maintenance of attention, subjects were given the task of reporting the direction change of the fixation point using a fibre-optic button box.

The cortical representation of retinotopic space was determined using a phase-encoded design in which the cardinal axes of visual space (‘polar angle’ and ‘eccentricity’) were mapped separately (Engel *et al.*, 1997). The stimuli consisted of two different high-contrast, multicolored checkerboard patterns, on a grey background, subtending a maximum visual angle of 24° × 24° (33.94° in diagonal). The ‘polar’ rotating wedge stimulus swept through the polar angle dimension clockwise or counter-clockwise, while the ‘eccentricity’ ring stimulus mapped eccentricity by starting from the centre of the visual field and expanding outward, or starting from the outer edge of the visual field and contracting inward. The ring and the wedge stimuli consisted of checkerboard patterns where brightness and colors changed at a rate of 8 Hz to maximize the neural responses of the visual areas of interest. The eccentricity stimulus traversed space with a logarithmically increasing rate as has been used previously (Sereno *et al.*, 1995; Conner *et al.*, 2004), whereas the arc angle of wedge stimulus was constant at 10°. These phase-encoded stimuli used a 64 s cycle, completing eight cycles per scan. Each scan was 512 s long, with 256 time points per functional scan volume. Four scans of this type were administered, two for eccentricity (1 expanding scan and 1 contracting scan) and two for polar angle (1 clockwise scan and 1 counter-clockwise scan). Paired clockwise/counter-clockwise and expansion/contraction scans were used in order to cancel the effects of residual hemodynamic phase delays.

The functional analysis of retinotopic data was completed using the FS-FAST software tools available in FreeSurfer software at (http://www.nmr.mgh.harvard.edu/freesurfer) (Dale *et al.*, 1999; Fischl *et al.*, 1999; Fischl *et al.*, 2001). Before statistical analysis, raw MR images were first motion-corrected to the midpoint volume (i.e. the 128th volume of 256 volumes for ‘polar’ and ‘eccentricity’ scans) of the first run using an iterated linearized weighted least squares method through the FS-FAST implementation of the AFNI 3dvolreg algorithm (Cox & Jesmanowicz, 1999). This algorithm provides a sum-of-squares estimate of average head motion throughout the fMRI scanning session, which was similar for both subject groups. The MR volumes were subsequently intensity normalized using the average in-brain voxel intensity.

Fast Fourier transform analysis was conducted on the time series of each voxel to statistically correlate retinotopic stimulus location with visual cortical anatomy. This analysis rejected low frequencies due to head motion or baseline drift and extracted both a magnitude and phase relative to the stimulus cycle frequency. Signal magnitude reflects retinotopic specificity, which can be low due to either lack of visually induced response or equivalent response to all retinotopic locations. The phase component of the signal codes specific retinotopic location.

### 2.4 Region of Interest Definition

Surface reconstructions of each subject's cerebral cortex were generated from three separate acquisitions of high-resolution anatomical images using the FreeSurfer version 5.3 software package (Dale *et al.*, 1999; Fischl *et al.*, 1999; Fischl *et al.*, 2001). The computed average cortical surface representation was inflated and then flattened by introducing a series of cuts to the 3-D surface to isolate the occipital pole (Sereno *et al.*, 1995). The resulting occipital patch was used to display the phases of the eccentricity and, separately, polar angle obtained from the retinotopy analysis of FS-FAST.

Visual areas V1, V2, and V3 were retinotopically identified on the occipital patch according to the criteria of Larsson and Heeger (2006). We drew the borders at the horizontal meridians to delineate V1-V2 and the outer edge of V3 and at the vertical meridian to delineate V2-V3. The visual areas were defined as labels on the surface. The labels of visual area, eccentricity, and polar angle, which were all defined on the surface, were projected into the anatomical voxels using FreeSurfer’s ‘mri_surf2vol’ command. We divided the visual areas into subregions, or regions of interest (ROIs), according to a set of criteria outlined in detail in Dawson *et al.*, (2013). First, we included only voxels that were within ±42.5° in polar angle of the oblique meridian in each quadrant of the visual field. This leaves a minimum 5° gap between ROIs in adjacent visual areas in order to minimize mixing of signal between different visual areas. Second, we included voxels that were significantly activated (p<0.05) in both eccentricity and polar angle runs. Third, we divided each visual area of each quadrant into three eccentricity bins: 0.25°-3.2° (central, e1), 3.2°-7.27 (mid, e2), 7.27-14° (peripheral, e3). For each ROI, the BOLD fMRI resting state time courses of all voxels within that ROI that passed the inclusion criteria were averaged together and normalized. For our network analyses, we divided the visual cortex into four quadrants based on hemisphere and dorsal/ventral position. Overall, we created 9 ROIs (3 visual areas x 3 eccentricity bins) in each of our 4 quadrants for a total of 36 ROIs per subject.

### 2.5 Statistics

Functional connectivity was assessed by calculating the Pearson correlation (Corr) and partial correlation (Pcorr) coefficients between all pairs of ROI resting state time courses within a particular quadrant, resulting in 36 Corr or Pcorr coefficients per quadrant per run per subject. Partial correlation refers to the correlation between two ROIs’ time-courses, from which the time courses of all other 7 ROIs in the network were regressed out.

Fisher’s z-transformation was applied on the Corr and Pcorr coefficients using the following:

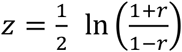

Fisher’s z-transformation is a variance-stabilizing transformation that converts Pearson’s r to the normally distributed z variable for use in statistical testing. A two-tailed one-sample t-test was conducted to assess whether Fisher transformed Corr/Pcorr coefficients were significantly different from zero in controls and separately in amblyopes. Next, a two-tailed two-sample t-test was conducted to compare Fisher transformed Corr/Pcorr coefficients between controls and amblyopes. Each subject had 8 Corr and 8 Pcorr values (4 quadrants * 2 runs) for each pair of ROIs. Using an alpha of 0.05, p-values were assessed for significance following False Discovery Rate (FDR) correction (Benjamini & Hochberg, 1995).

## 3 Results

We present functional connectivity data (Pcorr and Corr) separately for Amblyopes (Fig. 1A-C) and Controls (Fig. 1D-F). In Figs. 1A and 1D we show Pcorr and Corr coefficients for each pair of ROIs, averaged across each quadrant, run, and subject for Amblyopes and Controls, respectively. The Fisher transformed values are presented in Figs. 1B and 1E. Note that the results of Pcorr are presented in the bottom left portion of each matrix below the diagonal line, while the results of Corr are presented in the upper right portion.

**Fig. 1.**
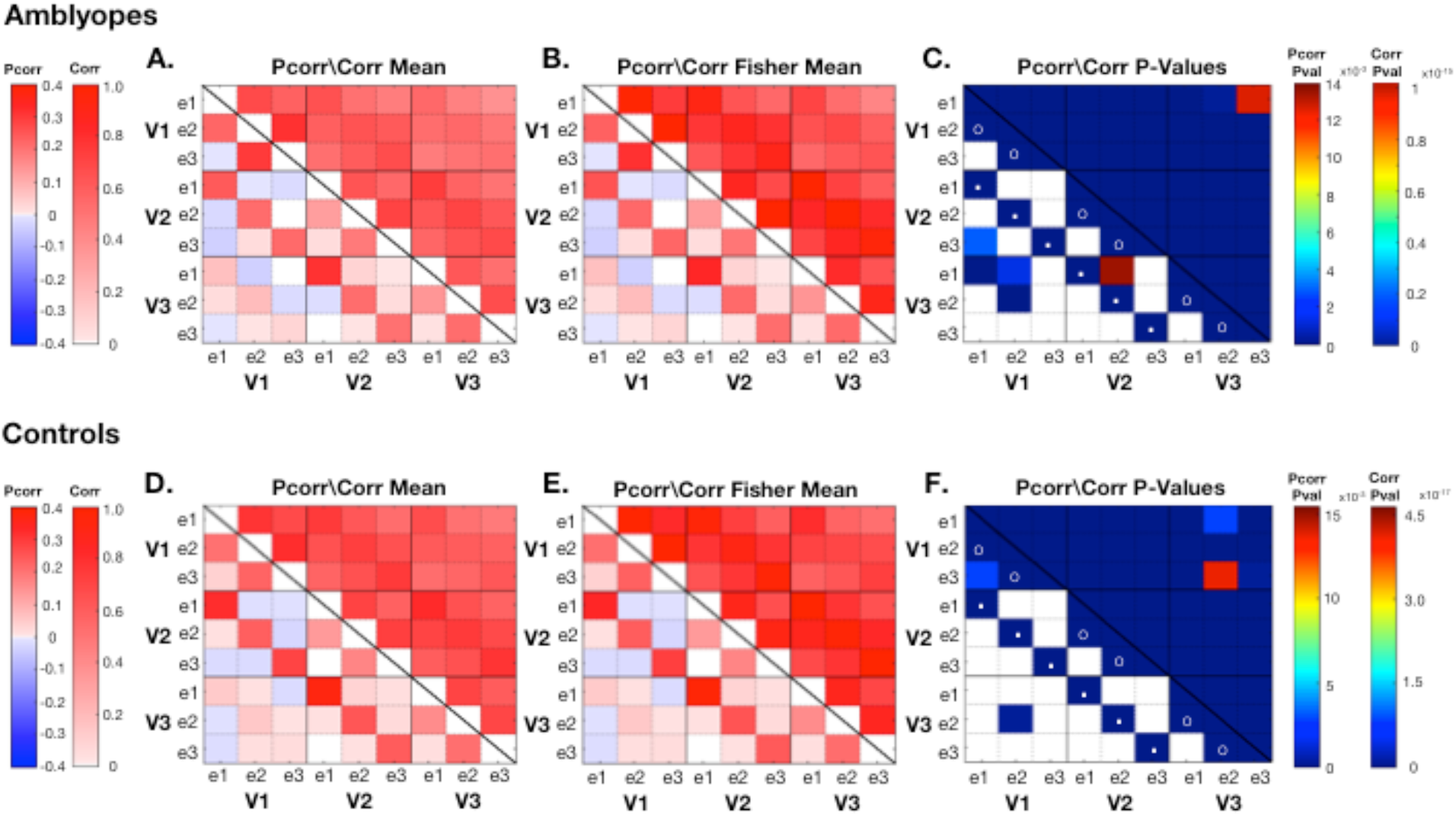
Matrices of mean Pcorr/Corr coefficients, Fisher transformed Pcorr/Corr coefficients, and *p*-values of two-tailed one-sample t-test of Fisher transformed Pcorr/Corr coefficients against zero for Amblyopes (A-C) and Controls (D-F). In (C), the FDR adjusted p-value threshold for Pcorr and Corr are 0.0139 and 1.06x10^-15^, respectively. In (F), the FDR adjusted p-value threshold for Pcorr and Corr are 0.0155 and 4.62x10^-17^, respectively. The p-value color bars represent the range of p-values from zero to the respective FDR adjusted p-value threshold for Pcorr and Corr. All Corr comparisons were significant in both Amblyopes and Controls. Pcorr comparisons coded in white in (C) & (F) are non-significant. Matrices are configured such that values for Pcorr are displayed on the bottom left portion of the matrix while values for Corr are displayed on the upper right.

In agreement with our previous findings in Dawson *et al.*, (2016), we observe retinotopic organization of functional connectivity within a quadrant in Controls as well as Amblyopes. There is an evident eccentricity effect: connections between adjacent eccentricity regions (e1-e2 and e2-e3) within an area [shown with open circles in Fig. 1C, 1F] had higher Pcorr and Corr than distant eccentricity regions (e1-e3). This was consistently seen in all visual areas in both subject groups. In addition, Pcorr revealed that connections between adjacent visual areas with same eccentricity (e.g., V1e1 & V2e1) [shown with dots in Fig. 1C, 1F] had higher connectivity measures than connections between distant visual areas and/or different eccentricity. However, this effect was not obvious in Corr and only manifested in Pcorr. Again, this was consistent in both subject groups.

In order to determine which connections were significant in Amblyopes and Controls, we conducted two-tailed one-sample t-tests on Fisher transformed Pcorr and Corr coefficients against zero. Fig. 1C and 1F show the results of the t-test for Pcorr and Corr for Amblyopes and for Controls, respectively, with an alpha of 0.05. For both Amblyopes and Controls, *all* Corr coefficients were significantly greater than zero (FDR corrected alpha of 1.06x10^-15^ and 4.619x10^-17^, respectively). Pcorr was more selective, however. Specifically, within-area, adjacent-eccentricity connections were significant, as were between-area, same-eccentricity connections in Amblyopes and Controls (FDR corrected alpha of 0.0139 and 0.0155, respectively). Comparisons coded in white did not meet statistical significance.

To test whether the functional connectivity was different between amblyopic patients and control subjects, in Fig. 2A we present the difference in the mean Pcorr/Corr between Amblyopes and Controls (Amblyopes minus Controls). Fig. 2B shows the difference between the Fisher transformed Pcorr/Corr values. We next conducted a two-tailed two-sample t-test comparing Fisher transformed Pcorr/Corr values between Amblyopes and Controls with alpha of 0.05. In each group, each subject contributed 8 Pcorr/Corr values per comparison (4 quadrants x 2 runs), which were pooled together and submitted to the two-sample t-test. Fig. 2C shows the t-statistic of the t-test and Fig. 2D shows the pooled standard deviation. The uncorrected p-values of the t-test are displayed in Fig. 2E. We found no significant difference in Fisher transformed Corr values between Amblyopes and Controls. Pcorr, on the other hand, revealed a few differences. A number of comparisons, all involving V1, yielded significantly different Fisher transformed Pcorr values. However, after correction for multiple comparisons, only the V1e2-V1e3 comparison remained statistically significant (Amblyopes > Controls, FDR corrected alpha of 0.0004) (Fig. 3).

**Fig. 2.**
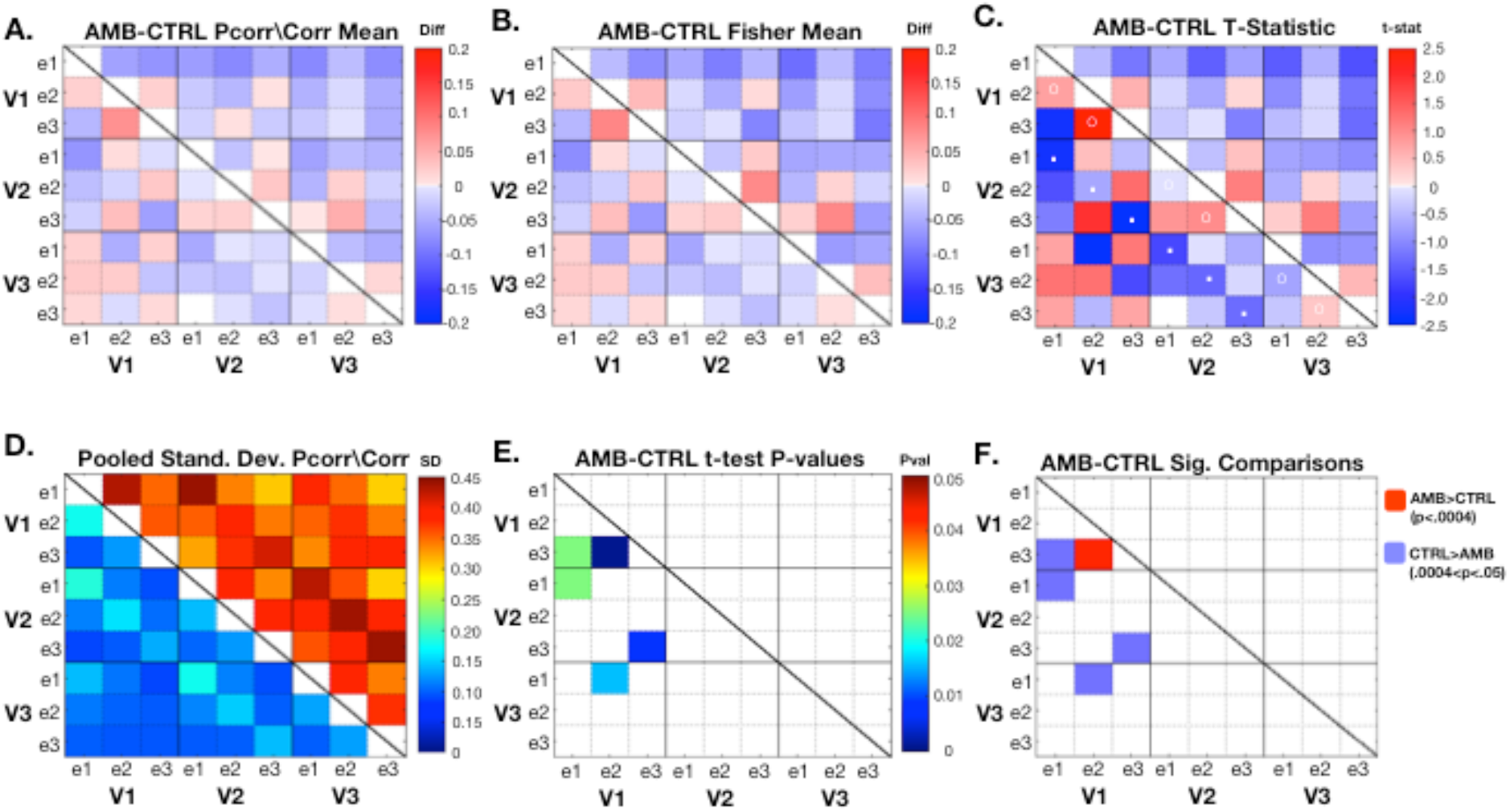
Matrices displaying differences between Amblyopes and Controls (AMB minus CTRL) in (A) Pcorr and Corr and (B) Fisher transformed Pcorr and Corr. (C) shows the t-statistic after performing a two-sample t-test on the Fisher transformed Pcorr and Corr values of Amblyopes vs Controls. (D) is a matrix displaying the pooled standard deviations of the Fisher transformed Pcorr and Corr values. (E) shows the resulting p-values from the t-test for comparisons that had p<0.05. In (F) we show which comparisons were significant after multiple comparisons correction and which were approaching significance. After FDR correction, the adjusted alpha became *p* = 0.0004, leaving only the comparison of the V1e3 - V1e2 connection significant (in dark red). Matrices are configured such that values for Pcorr are displayed on the bottom left portion of the matrix while values for Corr are displayed on the upper right.

**Fig. 3.**
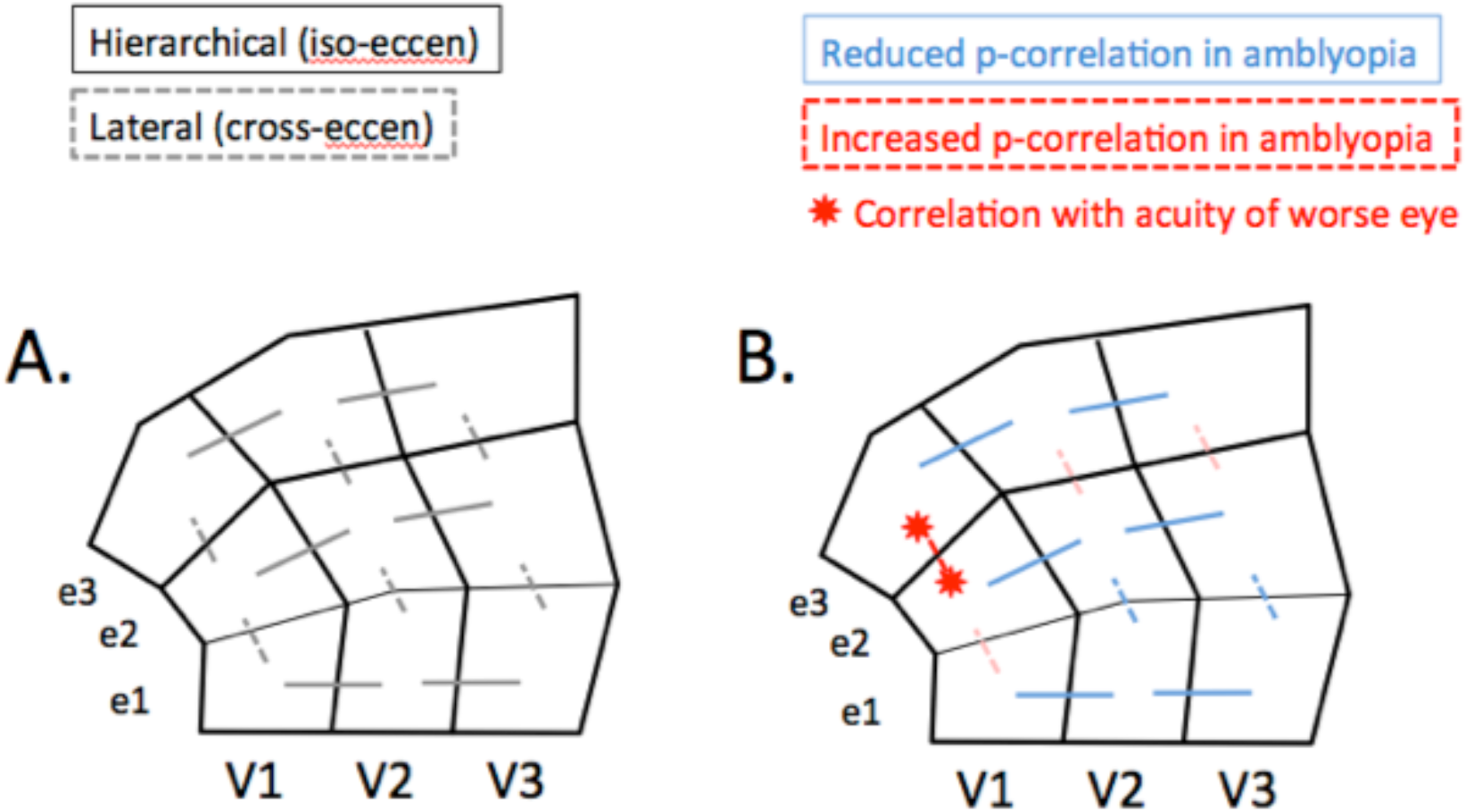
A schematic diagram for visualization of the pattern of differences between Amblyopes and Controls (AMB minus CTRL). The one partial correlation that survived correction for multiple comparisons is shown in red between the e3 and e2 eccentricity ROIs of V1. This correlation was stronger in amblyopes, and it correlated with worse eye acuity. It was also noted that similar trends existed (for both partial correlation and correlation) for enhanced within-area connectivity between e2 and e3 in V2 and V3 (shown in light red). Finally, the pervasive trend (both partial and regular correlation) for iso-eccentricity hierarchical connections between visual areas to be reduced in amblyopes compared to control is indicated in light blue.

In order to determine possible functional relevance of the above pair-wise Pcorr values we then tested for any association between this V1e2-V1e3 Pcorr values and our behavioral scores for worse eye acuity and for residual depth perception (stereoacuity). For the V1e2-V1e3 connection, low / high partial correlation was associated with more / less impairment of acuity. Inspection of this association indicated two potential subgroups rather than a continuum (Fig. 4A). To assess this, we performed K-means clustering with 2-7 clusters and plotted the within group sum of squares as a function of the number of clusters (Fig. 4B). Based on the point where this plot had its maximal curvature (the ‘elbow’ criterion), the cluster analysis identified two clusters: one with poor acuity scores and high partial correlation and another with better acuity scores and low partial correlation. In the case of stereo-acuity, however, no significant relationship was found. We also computed correlations with behaviour for the four connections that did not survive FDR correction (shown in light blue in Fig. 2F), but none were significant.

**Fig. 4.**
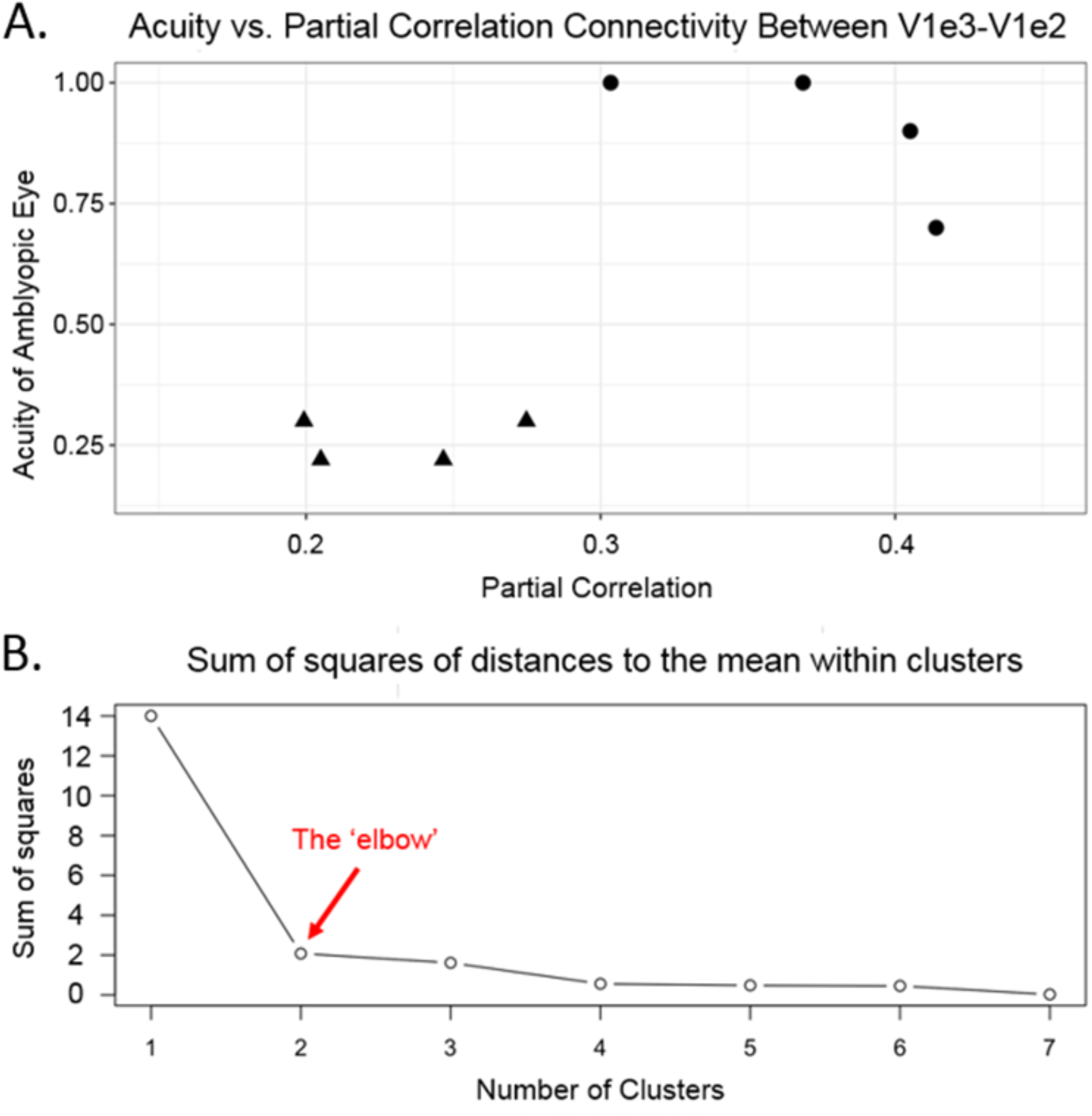
(A) Plot of V1e2-V1e3 partial correlation score versus acuity score of worse eye for all eight amblyopic subjects. The Pcorr assigned to the connection between V1e2 and V1e3 showed the only significant difference between amblyopic patients and control subjects (Fig. 2). The individual acuity scores are also given in Table 1. More impaired acuity is associated with higher Pcorr scores. The distribution of our sample suggests two subgroups, rather than a continuum or a spurious correlation driven by one outlier. (B) Estimation of the number of clusters within the scatter of data points presented in panel A. The estimaton is based on the K-means clustering sum of squares of distances to the mean within clusters versus cluster number. The cluster analysis identified two clusters. The “elbow” is the point of maximal curvature from two to seven clusters, indicating that two clusters account optimally for the variability of the data. One cluster has worse acuity scores and high partial correlation scores while the other has better acuity scores and lower partial correlation.

Two final analyses were conducted in response to observations we made regarding the pattern of differences between the amblyopic and control subjects seen in Fig. 2C. Recall that the bottom left portion of the matrix shows Pcorr, while the results of Corr are presented in the upper right portion. As can be seen in Fig. 2C, the six partial correlation connectivity measures between iso-eccentricity regions of adjacent areas (V1 and V2, and separately, V2 and V3) were reduced for amblyopic subjects [marked with white dots]. Testing with repeated measures the partial correlation across the six identified connections and subjects showed that the mean partial correlation was lower in Amblyopes versus Controls, t(88) = -2.6950, *p* = 0.0084.

Finally, it is also notable that for most pair-wise connections Corr is smaller in amblyopic than control subjects (upper right of matrix). We again applied repeated measures (with 36 connections/subject) statistics, and found that there was indeed a significant difference in correlation between Amblyopes and Controls, t(538) = -2.2866, *p* = 0.0226. This indicates that across the entire network, Amblyopes show decreased correlation (Corr) compared to Controls.

## 4 Discussion

We report a localized increase in *partial* correlation between regions within V1 as well as an overall reduction in correlation between iso-eccentric regions of V1 to V2 and V2 to V3 for subjects with amblyopia. This is the first study to analyze connectivity within and between ROIs that are smaller than an entire visual area in amblyopia. Our results first suggest cortical adaptions that cause a loss of effective resolution of visual representations in V1. Secondly, we find a general loss of normal drive from V1 to V2, and again from V2 to V3 that might build upon any reduced subcortical gain. Prior psychophysical evidence already exists in support of the idea that amblyopia results in these local and global perturbations of normal cortical connections. Our findings provide possible mechanisms for such psychophysical findings. Moreover, the significant association between more or less impairment of amblyopic eye acuity and increased partial correlation within V1, argues for the functional relevance of these findings.

### 4.1 Visual Cortex Organization in Amblyopia

In the current study, we found one connection with a significantly enhanced partial correlation between adjacent regions of interest in peripheral V1 in amblyopic subjects. The spatial layout of this increased connection is interesting considering that non-foveal representations (3-14 deg) were affected. A possible explanation could be decreased resolution in the vision of amblyopes. A shift toward a cruder retinotopy (i.e., larger receptive fields) could lead to an increased connectivity between these cross-eccentricity representations. One detailed study recently found that while cortical magnification is normal in the foveal field of strabismic amblyopes, the population-level receptive field sizes are enlarged for the amblyopic eye (Clavagnier *et al.*, 2015). Consistently, we found this increased Pcorr within V1 is associated with poor acuity in strabismics. Potentially similar associations have already been reported in normal subjects, although in this case between Vernier acuity and cortical magnification factor (Duncan & Boynton, 2003). Finally, McKee, Levi, and Movshon (2003) have found that strabismic subjects, especially those with very poor stereoacuity, tend to show poor monocular *acuity* yet relatively good monocular *contrast sensitivity*. These authors suggest that this pattern of psychophysical performance could result if V1 in strabismic subjects contains a greater number of monocular neurons (possibly with larger receptive fields). The trend we found for poor stereo-acuity in subjects with poor acuity is consistent with this model.

In addition to the shorter-range connectivity differences within V1, our results also include longer-range connectivity differences within the V1-V3 visual cluster. As might be expected from the generally reduced fMRI responses during task-based fMRI, overall connectivity was reduced in amblyopia compared to controls. These reduced longer-range connections seen in amblyopia may be representative of reduced feedforward connections within this hierarchical visual system (Li et al., 2011). Thus, deficits may increase as information goes up the hierarchy. Consistently, Muckli *et al.*, (2006) showed that responses to amblyopic eye stimulation were progressively reduced in higher-tier cortical areas that emphasize central vision. This concept can serve as a plausible explanation for altered temporal and parietooccipital cortical activity observed in amblyopes, as well as other behavioral deficits (Levi, 2006), such as impaired visuomotor behavior (Niechwiej-Szwedo *et al.*, 2012, 2014) or depth perception (Joly & Frankò, 2014).

One possible source of reduced feedforward connections is inter-ocular suppression. Behaviorally, this has been found to encompass the central 20° of the visual field, although the greatest impact is in the central-most regions (Babu *et al.*, 2013; Hess *et al.*, 2014; Babu *et al.*, 2017). Visual evoked potentials (VEP) and fMRI and have demonstrated that amblyopic eye stimulation generally results in lower activation in V1, compared to the fellow eye or compared to control subjects (e.g., Barnes, *et al.*, 2001; Lee *et al.*, 2001; Lerner *et al.*, 2006; Muckli *et al.*, 2006; Conner *et al.*, 2007b; Hess *et al.*, 2009a; Wang *et al.*, 2012). Whether such physiological attenuation or suppression grows in strength or simply “passes-through” subsequent higher-tier areas is not known (Levi, 2006; Jurcoane *et al.*, 2009; Chen & Tarczy-Hornoch, 2011). Suggestive evidence for active suppression of the foveal representations, including those beyond the occipital pole in the occipito-parietal cortex (Wandell *et al.*, 2005), has also been reported when amblyopic eye stimulation with fellow eye open was contrasted with fellow eye closed (Conner *et al.*, 2007b). Additional studies may be beneficial in resolving the debate between reduced input gain and active suppression. This can be done by identifying decreased connectivity in amblyopia of “higher-tier” cortical regions, comparing the strength of signal reduction to that of earlier cortical regions, and determining the functional deficits associated with abnormalities in these higher-order regions.

For comparison, we mention here recent results from resting state studies in subjects with *complete* loss of vision in childhood (Bock *et al.*, 2015; Butt *et al.*, 2015). Results suggest that while the gross retinotopic structure of correlations from V1 to V2, and from V1 to V3 is maintained in early-blind subjects, *enhanced* correlations between cortical areas may exist. Therefore our finding of *reduced* correlations between adjacent visual areas would suggest a different mechanism in amblyopia. It seems likely that complete blindness makes reallocation of function in visual cortex possible, and V1 likely participates in other whole-brain networks (e.g., Amedi, *et al.*, 2003). For example, there is evidence from blind subjects that connectivity with regions as distant as Broca’s area in frontal cortex increases substantially (Sabbah *et al.*, 2016). This would contrast with amblyopia where a disconnection of V1, at least locally, might result from the discordant binocular signals and resultant suppression to avoid diplopia (e.g., Conner *et al*., 2007b).

In the future, it would be ideal to include pediatric subjects and potentially acquire data longitudinally. It has been reported recently that in a large sample of children with both types of amblyopia, perceptual suppression of the weak eye’s stimulus is common and testable as young as age 3 years (Birch et al, 2016). We know these children have highly asymmetrical interocular suppression that can vary in severity, but the specific neural mechanisms are still unidentified. Resting state studies are also feasible for very young children and offer the real potential to track etiological mechanisms and distinguish them from normal developmental changes. Ultimately, lagged-based measures of effective connectivity might allow us to further distinguish feedback connections from feedforward input. Finally, functional MRI at 7T enables smaller regions of interest that distinguish cortical columns or layers (Chaimow et al., 2017; Olman et al., 2017; Guidi et al., 2016; Huber et al., 2017; Yacoub et al., 2007). This can potentially support the differentiation of activity in input and output layers of areas in the visual cortex.

### 4.2 Previous Resting State Studies

The complex retinotopic organization of the human visual system is relatively well-known (e.g., Wang *et al.*, 2015), and the degree to which this information is evident in the spontaneous fluctuations in fMRI data has been an active topic of investigation. Others have also extracted BOLD signals from functionally defined areas (V1, V2, or V3) in normal subjects, and found that the resting state correlation pattern between areas reflects the widespread eccentricity organization of visual cortex, in which the highest correlations are observed for cortical sites with iso-eccentricity representations (Buckner & Yeo, 2014; Arcaro *et al.*, 2015). Previous rs-MRI of visual cortex at 7T also revealed retinotopically precise correlations between V1, V2, and V3 over the same and opposite hemisphere (Gravel *et al.*, 2014; Raemaekers *et al.*, 2014; Genç *et al.*, 2015). However, such functional connectivity between regions smaller than a cortical area was observed only after the global contributions from a large-scale bilateral network were removed (Raemaekers *et al.*, 2014). Another study that specifically included measures from subcortical nuclei also emphasized the mixture of independent signal sources in the correlation structure measured in visual cortex (de Zwart *et al.*, 2013). Moreover, in our previous study (Dawson *et al.*, 2016), we have explicitly shown that correlation-based functional connectivity is non-selectively high across lower visual areas V1-V3, even between regions within these areas that do not share direct anatomical connections. We therefore suggest that the mechanisms underlying correlation based functional connectivity likely involve network effects caused by the dense anatomical connectivity within this cortex and feedback projections from higher visual areas. In contrast, *partial* correlation, which minimizes network effects, follows expectations based on the direct anatomical connections verified in non-human primate visual cortex better than correlation. Our findings with partial correlation discussed above are thus based on well-validated methods.

As mentioned in the introduction, a few studies of larger scale rs-fMRI in subjects with amblyopia have been reported recently (Lin *et al.*, 2012; Ding *et al.*, 2013; Wang *et al.*, 2014; Liang *et al.*, 2016). Both increased and decreased correlation compared to normal subjects were observed. However, none of these studies compared correlation to partial correlation. We suspect that many of these findings may not be replicated with a more selective partial correlation test. Moreover, for these whole-brain measures, the interpretation of how the findings relate to functional consequence can be challenging. In only two instances, modest but suggestive correlations were found with some behavioral measures. Tan *et al.*, (2016) found an association between their ALFF measure (amplitude of low frequency fluctuation) in angular gyrus and duration of strabismus, while Liang *et al*., (2016) correlated the standardized ALFF of the precuneus to the amount of optical anisometropia (but not to acuity). Finally, the effective (lag-based) connectivity (Corr) loss from LGN to V1 reported by Li *et al.*, (2011) during visual stimulation of the worse eye correlated with the worse eye acuity (degree of amblyopia). Thus, the suggestive relationship of *partial* correlation with acuity that we found for short-range connections within V1 (Fig. 4) may derive in part from precortical alterations. That this local effect within resting V1 was only detectable with partial correlation highlights the emerging understanding of amblyopic deficits as manifestations of abnormal interactions between distinct neural populations that may or may not be physically adjacent. Finally, we point out that even the application of partial correlation analysis to previously published resting state data from amblyopic subjects could reduce the network effects and the effect of common input. We expect that this could provide a much more selective picture of amblyopic cortical adaptations than the current literature suggests.

### 4.3 Conclusions

Retinotopically guided functional connectivity analysis of areas V1, V2, and V3 shows reduced interaction between V1 and V2 and separately between V2 and V3 in adult amblyopia. In addition, only partial correlation selectively isolated abnormally strong functional connectivity across eccentricities within V1. This effect was strongest in strabismic subjects with the worst acuity. Future studies including more adults with anisometropia and children with both types of amblyopia would be very useful. It remains unknown when during visual cortex development resting state connectivity becomes abnormal in amblyopic subjects. Measurement of rs-fMRI connectivity, with a special emphasis on partial correlation, requires minimal assumptions about the adaptations that underlie amblyopia, and seems suitable to reveal additional neural substrates beyond those currently recognized. Finally, we stress that the use of partial correlation analysis is applicable to all resting state connectivity studies and may offer additional specificity and sensitivity for the characterization of a wide array of brain disorders.

## Abbreviations

(LGN): lateral geniculate nucleus
(rs-fMRI): resting state functional magnetic resonance imaging
(Corr): correlation
(Pcorr): partial correlation

## Acknowledgements

The authors wish to acknowledge Debra Dawson and Dr. Felix Carbonell for helpful discussions. We also thank Dr. Robert Hess for performing ophthalmic exams.

